# Toxic SOD1 trimers are off-pathway in the formation of amyloid-like fibrils in ALS

**DOI:** 10.1101/2021.08.17.456705

**Authors:** Brianna Hnath, Nikolay V. Dokholyan

## Abstract

Accumulation of insoluble amyloid fibrils is widely studied as a critical factor in the pathology of multiple neurodegenerative diseases, including amyotrophic lateral sclerosis (ALS), a fatal neurodegenerative disease. Misfolded Cu, Zn superoxide dismutase (SOD1) was the first protein linked to ALS, and non-native SOD1 trimeric oligomers were recently linked to cytotoxicity, while larger oligomers were protective to cells. The balance between trimers and larger aggregates in the process of SOD1 aggregation is, thus, a critical determinant of potential therapeutic approaches to treat ALS. Yet, it is unknown whether these trimeric oligomers are a necessary intermediate for larger aggregate formation or a distinct off-pathway species competing with fibril formation. Depending on the on- or off-pathway scenario of trimer formation, we expect drastically different therapeutic approaches. Here, we show that the toxic SOD1 trimer is an off-pathway intermediate competing with protective fibril formation. We design mutant SOD1 constructs that remain in a trimeric state (super stable trimers) and show that stabilizing the trimeric SOD1 prevents formation of fibrils *in vitro* and in a motor neuron like cell model (NSC-34). Using size exclusion chromatography we track the aggregation kinetics of purified SOD1 and show direct competition of trimeric SOD1 with larger oligomer and fibril formation. Finally, we show the trimer is structurally independent of both larger soluble oligomers and insoluble fibrils using circular dichroism spectroscopy and limited proteolysis.

**Significance Statement:** Protein aggregation into large insoluble species is a hallmark of many neurodegenerative diseases, but recent evidence suggests that smaller soluble aggregates are responsible for neuronal death. Depending on whether these toxic oligomers appear on- or off-pathway to larger aggregates, which is currently unknown, the strategies for pharmaceutic intervention would be drastically different. Here, we determine that stabilizing the trimeric form of SOD1 reduces larger aggregate formation while increasing toxicity to cells. Trimeric and larger aggregate concentrations have a strong negative correlation over time, and the structure of the trimer is significantly different from larger soluble and insoluble aggregates. Our findings show that formation of trimeric SOD1 is directly competing with that of larger aggregates.

## Introduction

Cu, Zn superoxide dismutase 1 (SOD1) mutations account for 15-30% of familial ALS (fALS) (family history) cases and 1-2% of sporadic (sALS) cases (1) (2). Recent studies have discovered misfolded SOD1 in sALS patient tissue samples without any genetic mutations(3) (4), and post translational modifications to SOD1(5) (6) (7) (8), either occurring naturally or from environmental toxins (9) (10) (11), have been shown to destabilize SOD1 leading to aggregation. Mutations (12) (13), loss of metals (14) (15), crowding (16), and post translational modifications to SOD1 lead to destabilization of the dimer interface. De-metallated SOD1 monomers are highly prone to aggregation (17) and the large insoluble inclusion bodies they form have been studied since the protein was linked to ALS (18) (19) (20).

The formation of large, insoluble, amyloid fibrils is thought to be a main factor of disease progression and cell death in a number of neurodegenerative disorders, such as Alzheimer’s disease, Parkinson’s disease, and amyotrophic lateral sclerosis (ALS) (21) (22) (23) (24). Despite new studies indicating that smaller soluble oligomers are likely to be toxic in many of these disorders, fibrils remain at the heart of the *amyloid cascade hypothesis* (25) (26), in which amyloid fibrils are the principal toxic species responsible for neurodegenerative diseases and remain diagnostic and therapeutic targets (27) (28) (29) (30). Recent studies of SOD1 aggregation paint a different picture from a traditional amyloid cascade hypothesis: Proctor et al. found that non-native SOD1 trimers (31) are toxic to motor neuron-like cells, while Zhu et al. found that larger aggregates are protective (32). We posit that protein aggregation is a protective mechanism against increased population of misfolded proteins. Most globular proteins are marginally stable and various triggers, such as genetic variations (12) (13), protein modifications (5) (11), oxidative stress (6), promote the population of misfolded proteins. Some of the states of these proteins may induce downstream toxicity, while aggregation serves a protective role by shifting the population of toxic species towards potentially inert amyloid fibrils. This scenario is opposite to the dominating amyloid cascade hypothesis where the amyloid fibrils themselves are believed to be toxic. If aggregation is favorable for cell survival, we expect that the aggregation process would compete with the formation of toxic oligomers. Zhu et al.(32) observed the premise for this hypothesis in experiments when he designed SOD1 mutants that promoted large aggregate formation but destabilized toxic trimers. These mutant SOD1 species did not impact cell viability even in the presence of large amounts of aggregates.

How do SOD1 aggregates protect cells from toxic intermediates? There are two possible scenarios: SOD1 trimers appear (i) on-pathway or (ii) off-pathway to aggregation. In the former case, rate of the large aggregate formation may significantly exceed the lifetime of the trimer, while in the latter case, large aggregates compete with the trimeric intermediates. Understanding which scenario happens in cells is critical for developing pharmaceutical strategies: If trimeric SOD1 appears off-pathway, therapeutic strategies should focus on promoting fibril formation instead of breaking apart fibrils (Figure 1). If trimeric SOD1 is on-pathway, we expect arresting the protein aggregation process to inhibit formation of trimers.

**Fig. 1.**
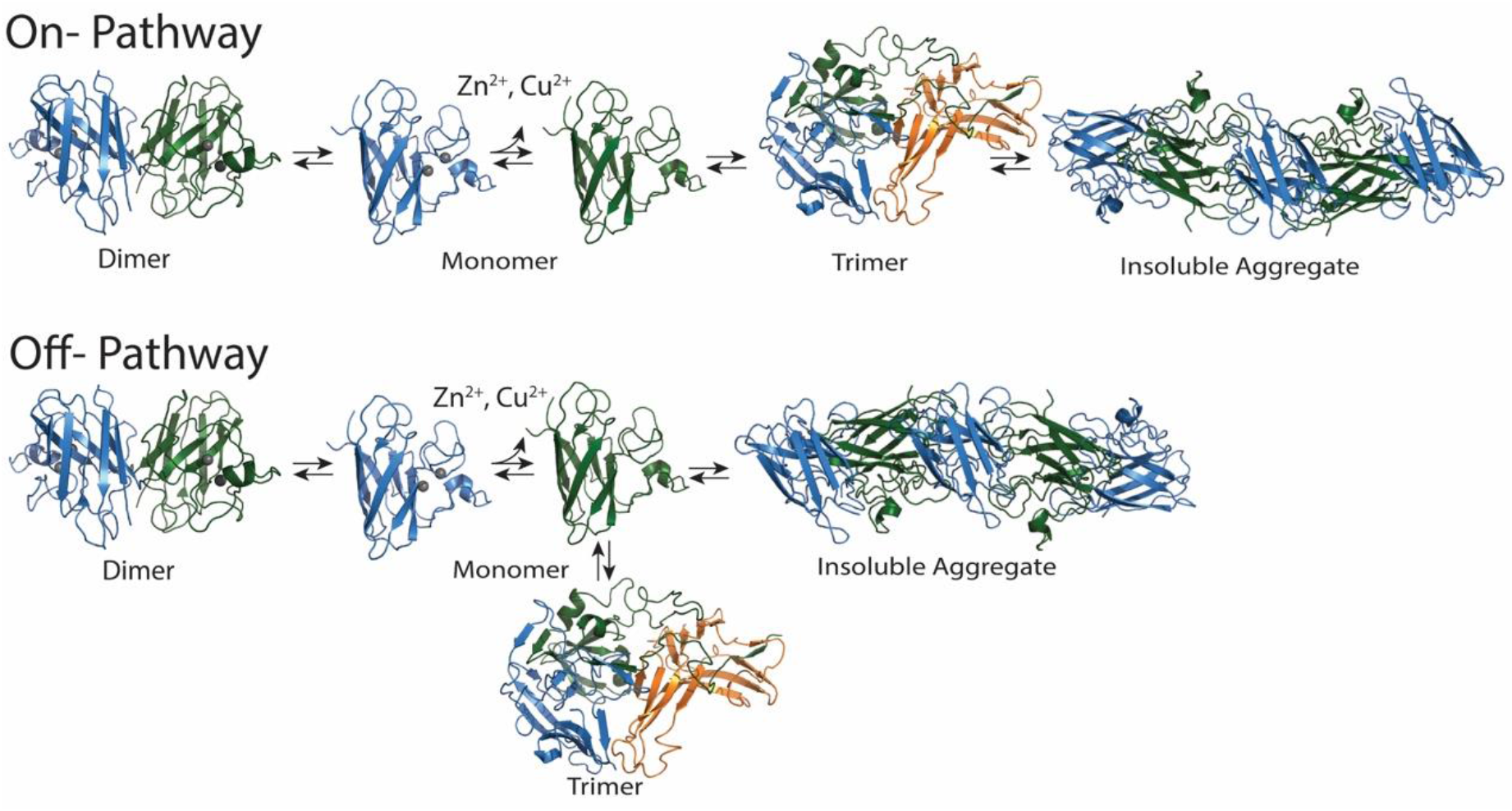
The formation of soluble intermediates for SOD1 is thought to happen either on- or off-pathway. On-pathway the formation of the trimer is a necessary step in the formation of larger insoluble fibrils. Off-pathway the formation of the toxic soluble trimer directly competes with insoluble fibril formation.

Here, we show that trimeric SOD1 is off-pathway in the formation of protective SOD1 inclusion bodies. We combine point mutations to stabilize the trimer which shows reduced inclusion body and amyloid-like aggregate formation in a Thioflavin T (ThT) amyloid aggregation study and in motor neuron like NSC-34 cells. The aggregation kinetics display a strong negative correlation between trimer and aggregate concentrations further suggesting that trimers are an off-pathway state. Finally, trimeric SOD1 is structurally different from both native dimeric form and larger soluble and insoluble aggregates. These results explain why SOD1 inclusion body and large aggregate formation is protective to cells and show that SOD1 trimers are structurally distinct from larger aggregates and native dimers. Finding that SOD1 trimers are formed off-pathway suggests two possible therapeutic intervention strategies: (i) inhibit native dimer dissociation, which results in loss of metals and misfolding, and/or (ii) promote SOD1 aggregation, thereby depleting the formation of SOD1 trimers.

## Results

In order to regulate the population of SOD1 trimers, we perform thermodynamic stabilization of these states by combining in different pairings trimer stabilizing mutations, identified by Proctor et al. (31). We calculate the effect of the mutations on the free energy of the dimer and different proposed trimer structures using the molecular design suite Eris (33) (34). The most promising combinations to stabilize the trimer are overexpressed and purified from yeast cells. After purification, the proportion of trimer to other species and the stability of the trimer over time are both tested using native PAGE and size exclusion chromatography (SEC). De-metallated F20L-H46Q SOD1 and H46Q-G108H SOD1 both have a large proportion of trimers directly after purification. These purified trimers remain in this state: their population does not significantly change over a 48 hour time course (incubate at 37°C in 20 mM Tris 150 mM NaCl pH 7.4, use SEC to track changes) (Supplementary Figure 1). Stabilizing the trimeric form allows us to investigate the structure of the more commonly transient species.

### Stabilizing the trimer reduces inclusion body and amyloid-like fibril formation

Fibrillization assays using Thioflavin T (ThT) to track the formation of β-sheet rich fibrils are widely used to study aggregation of several different proteins (35) (36) (37). Studies comparing the fibrillization of different ALS occurring SOD1 mutations have already seen a reduction in fibrillization propensity for some deadly mutants such as E100G, G37R, and H46R (38). If the SOD1 trimer is an intermediate in the formation of fibrils, mutants stabilizing trimers should not affect the formation of fibrils. We perform a fibrillization assay using SOD1 WT, A4V, F20L, and two of our super-stable trimer mutants. A4V has an increase in fibrillization compared to WT while F20L reduces fibrillization and the super stable trimers do not form any fibrils (Figure 2A).

**Fig. 2.**
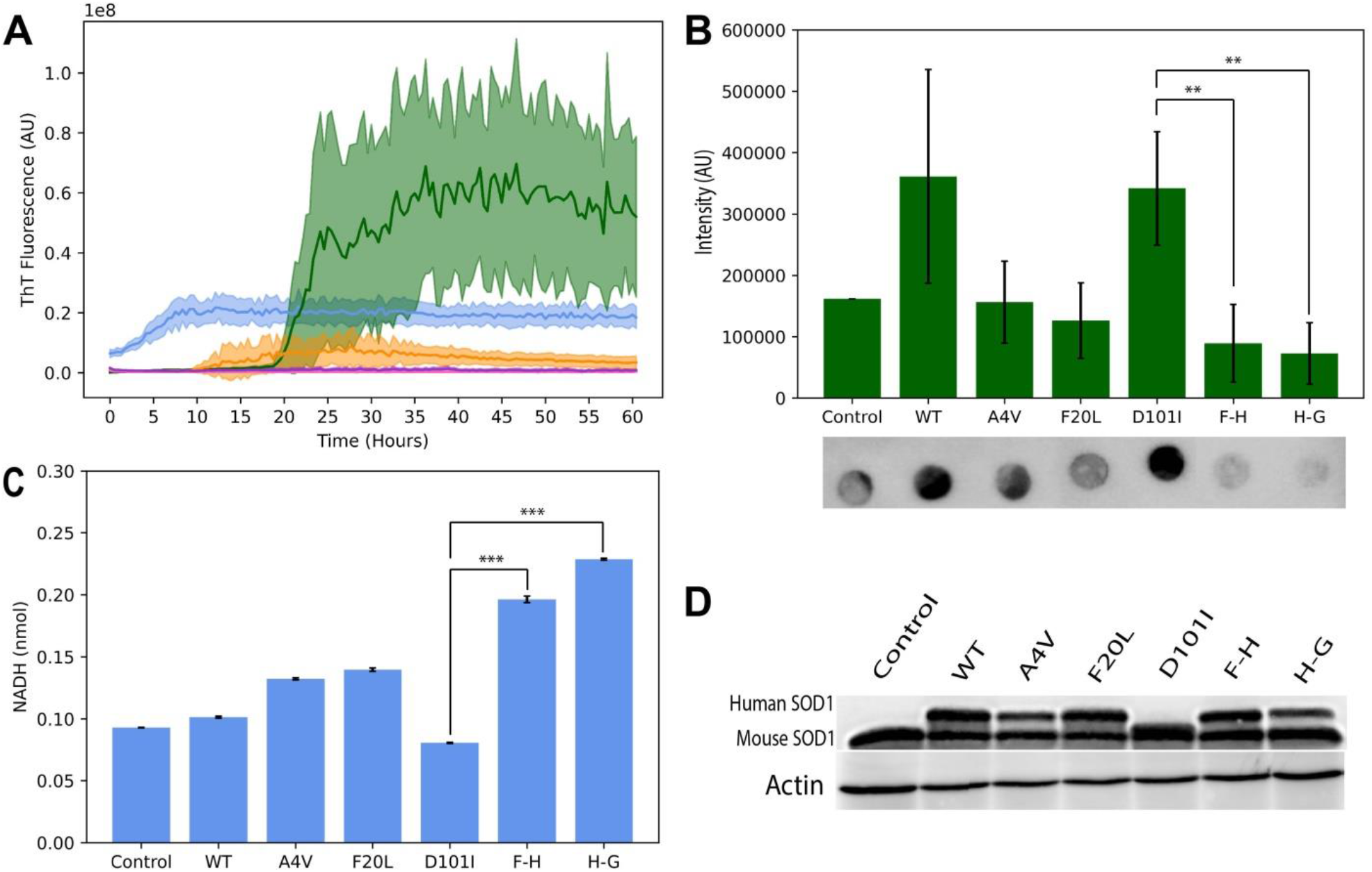
Stabilizing the trimeric form of SOD1 reduces insoluble fibril formation and increases cell death. (A) ThT expression shows a large amount of amyloid aggregation in A4V mutants (green) compared to WT (blue) and this expression reduces as the trimer is stabilized with slight stabilization by F20L (orange) and further stabilization by F20L-H46Q and H46Q-G108H (pink and purple). (B) Dot blots of the insoluble protein fraction of NSC-34 cells expressing several SOD1 mutants show a larger portion of insoluble SOD1 in WT, A4V, and trimer destabilizing mutant D101I compared to trimer stabilizing mutants (F20L-H46Q (F-H) and H46Q-G108H (H-G)) showing that stabilizing the trimer reduces inclusion formation. (C) NADH (lactate dehydrogenase assay) intensity shows a significant increase in cell death when trimers are stabilized (F20L, F20L-H46Q (F-H), and H46G-G108H (H-G)) and a decrease in cell death when trimers are destabilized (D101I) compared to native mutant A4V. (D) Expression of the different mutants of human SOD1, D101I has a lower molecular weight human SOD1 band than all other SOD1 constructs but 2 bands (human and mouse) are still visible and other human specific antibodies still identified D101I. (** p<5×10^−2^) (*** p<10^−4^)

A similar phenotype is shown in NSC-34 cells transfected with the same SOD1 mutants along with D101I, which destabilizes the trimer. Three days post transfection the cells expressing trimer stabilizing mutants (Supplementary figure 2) have a significant decrease in the amount of inclusions (visualized by anti-SOD1 dot blots of the insoluble protein fraction), while D101I has even more insoluble SOD1 than A4V (Figure 2B). The toxicity of the stabilized trimers is similar to what was observed by both Proctor et al. and Zhu et al. (31) (32), stabilizing the trimer (F20L, F20L-H46Q, and H46Q-G108H) increases cell death while destabilizing the trimer (D101I) rescues the cells when compared to WT and A4V using a lactate dehydrogenase activity assay to evaluate cell viability (Figure 2C). These results confirm that larger SOD1 aggregates are protective to cells and that SOD1 trimers are directly competing with fibril formation.

### Trimer population negatively correlates with aggregate population

Many models for the aggregation of proteins focus on the nucleation and growth processes, but often make the assumption that aggregation is only a linear process (eg. Figure 1, on-pathway), similar to how crystal formation occurs (39) (40) (41). Validation of the models is primarily based on data from ThT aggregation experiments. ThT only binds across β-sheets in amyloid fibrils, thus, this method cannot account for protein conformations that have low percentages of β-sheets as, for example, in SOD1 trimers. Hirota and Hall developed a model for protein aggregation that removes the assumption that aggregation is a linear process (42) (43) and also includes changes in concentrations over time as an output for their model instead of equating the system to ThT curves. They show that during protein aggregation where the intermediate is on-pathway, the concentration of the intermediate has a positive correlation with the concentration of larger aggregates. The opposite occurs when the intermediate is off-pathway and competing with larger aggregate formation; in an off-pathway case there is a strong negative correlation between the intermediate and larger aggregates.

We test whether SOD1 trimers are on- or off-pathway by following the aggregation process of de-metallated trimer forming SOD1 mutants (A4V and F20L) for at least a week so the formation of different species at a set concentration (30 μM) is close to equilibrium. Khare et al previously demonstrated that obligatory steps for SOD1 aggregation are the dissociation of native dimers and the loss of metals (17). Hence, to follow aggregation process, we start with de-metallated samples so that only apo-monomers, trimers, and larger aggregates can form. We then perform SEC separation on aliquots taken every 0.5-2 hours for 3 days and quantify the concentration changes. The same amount of protein is loaded at each time point allowing us to account for fibril formation (fibrils are too large to quantify using SEC) by calculating any loss in total mass. All the samples have a significant, strong, negative correlation between trimers and larger aggregates/fibrils indicating that the trimers are off-pathway and competing with fibril formation (Table 1, Supplementary figures 3-7).

**Table 1.**
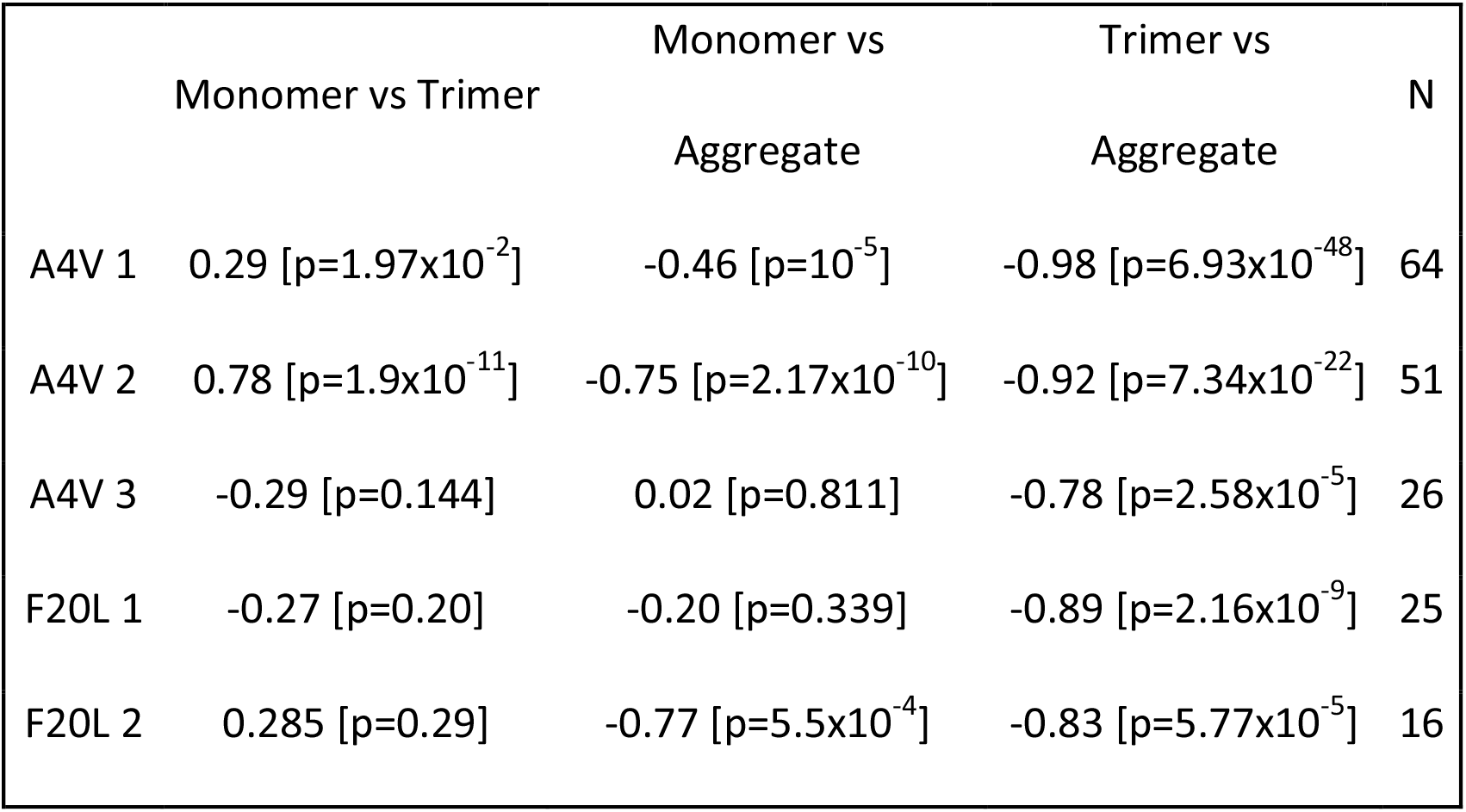
Size exclusion chromatography time courses show a significant strong negative correlation between trimers and aggregates which matches the competition between the two forms if the trimer is off-pathway.

### Trimeric SOD1 is structurally independent from larger oligomers and fibrils

We compare limited proteolysis of SOD1 trimers to that of large insoluble fibrils and find two exposed regions in the fibrils which are not exposed in the trimers suggesting there is a structural difference between these species (Figure 3, A). When WT and A4V insoluble fibrils, as well as F20L-H46Q and H46Q-G108H trimers, are digested with pepsin at four different time points (1, 5, 10, or 20 minutes) they feature five similar cut sites, three between residues 5-10 then at residue 116 and residue 144. The insoluble fibril samples have additional cut sites between residues 30-45 and residues 120-135, which do not appear in the trimer samples. Our findings suggest that those two regions are not exposed in the trimer samples and that the trimer is structurally independent from insoluble fibrils.

**Fig. 3.**
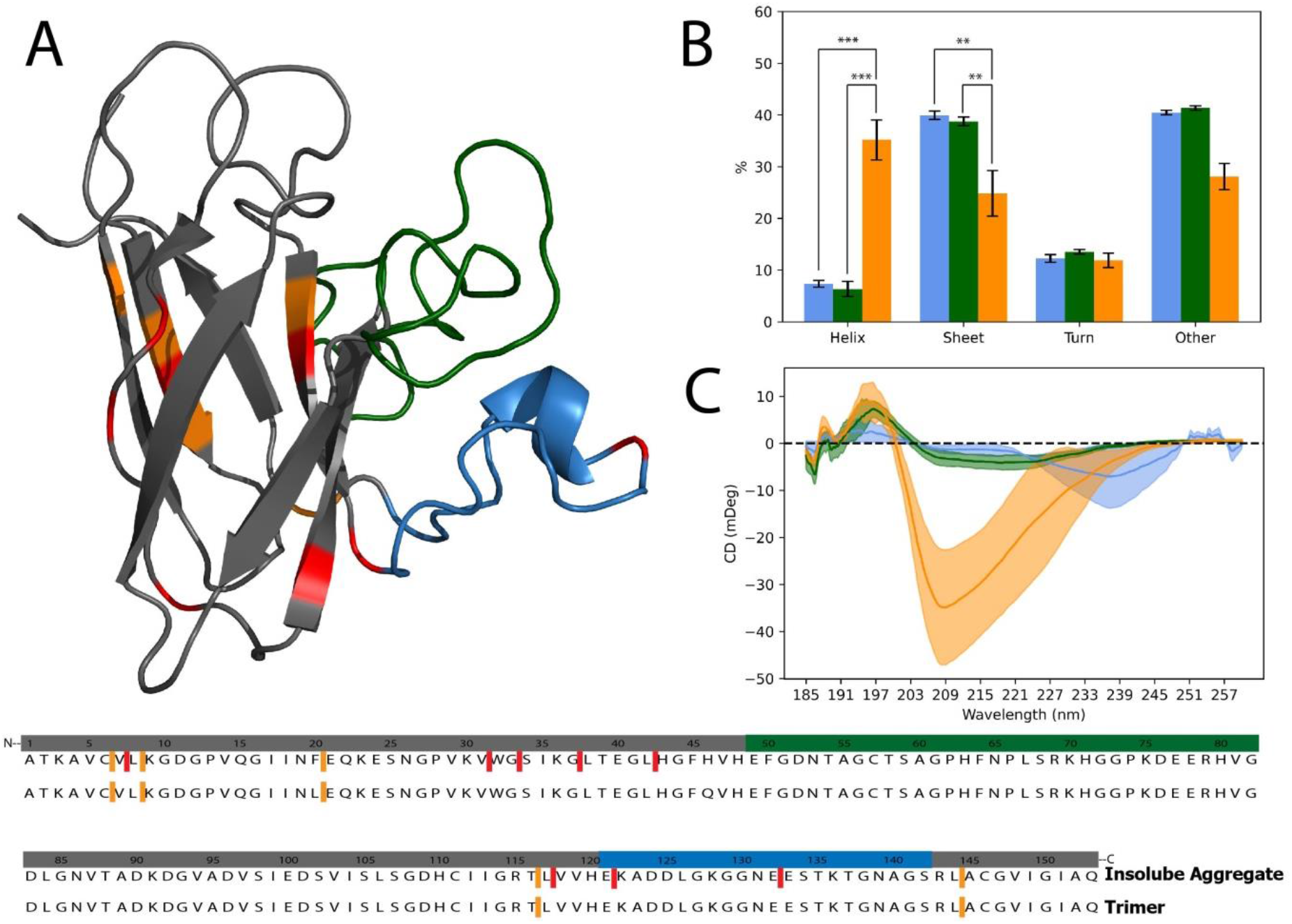
Trimeric SOD1 is structurally independent from monomeric and larger soluble and insoluble aggregates. (A) Pepsin digestion of two insoluble aggregate samples (WT and A4V) as well as two trimer samples (F20L-H46Q and H46Q-G108H) show five similar cut sites (orange) as well as eight additional cut sites (red) which are only present in the insoluble fibrils. (B) CD measurements show that SOD1 trimers (orange) have significantly more helical regions and less Beta sheet regions than monomers (blue) and larger soluble aggregates (green) (** p<5×10^−2^) (*** p<10^−4^).

We observe a similar structural difference between SOD1 trimers, monomers, and larger soluble oligomers. Circular dichroism spectroscopy (CD) of mutants and WT of SOD1 in each of the different conformations shows a significant increase in the percent of helix regions (monomer vs trimer *p* = 4×10^−4^, aggregate vs trimer *p* <10^−4^) and a significant decrease in the percent of β sheet regions (monomer vs trimer *p* = 2×10^−2^, aggregate vs trimer *p* = 8×10^−3^) between monomers/ large oligomers and trimers (Figure 3B). If the trimer is an intermediate species on the pathway of large fibril formation, we would expect it to have a similar structure to monomers and larger aggregates. Hence, structural analysis confirms that SOD1 trimers are off the aggregation pathway.

## Discussion

Protein aggregation is likely a natural, protective, cell response to stress, yet therapeutics are still being developed that aim to break down these protective aggregates (44) (45) (46) (47). Since many aggregating proteins linked to neurodegenerative disorders have been shown to cause a toxic gain of function, it is crucial to determine if these toxic conformations are structurally independent from the native form (48) (49) (50). Determining if these soluble oligomers are on- or off-pathway directly dictates the therapeutic strategy. For on-pathway intermediates we expect to use fibrils as a biomarker for cell health, and targeting the fibril might also prevent the toxic intermediates from forming. Alternatively, for off-pathway intermediates, promoting fibril formation could potentially rescue cells since the fibrils directly compete with the intermediates. Our study strongly suggests that trimeric SOD1 is off-pathway. We propose that accumulation of misfolded SOD1 triggers protective aggregation, which leads to the formation of both fibrils and toxic off-pathway trimers (Figure 4).

**Fig. 4.**
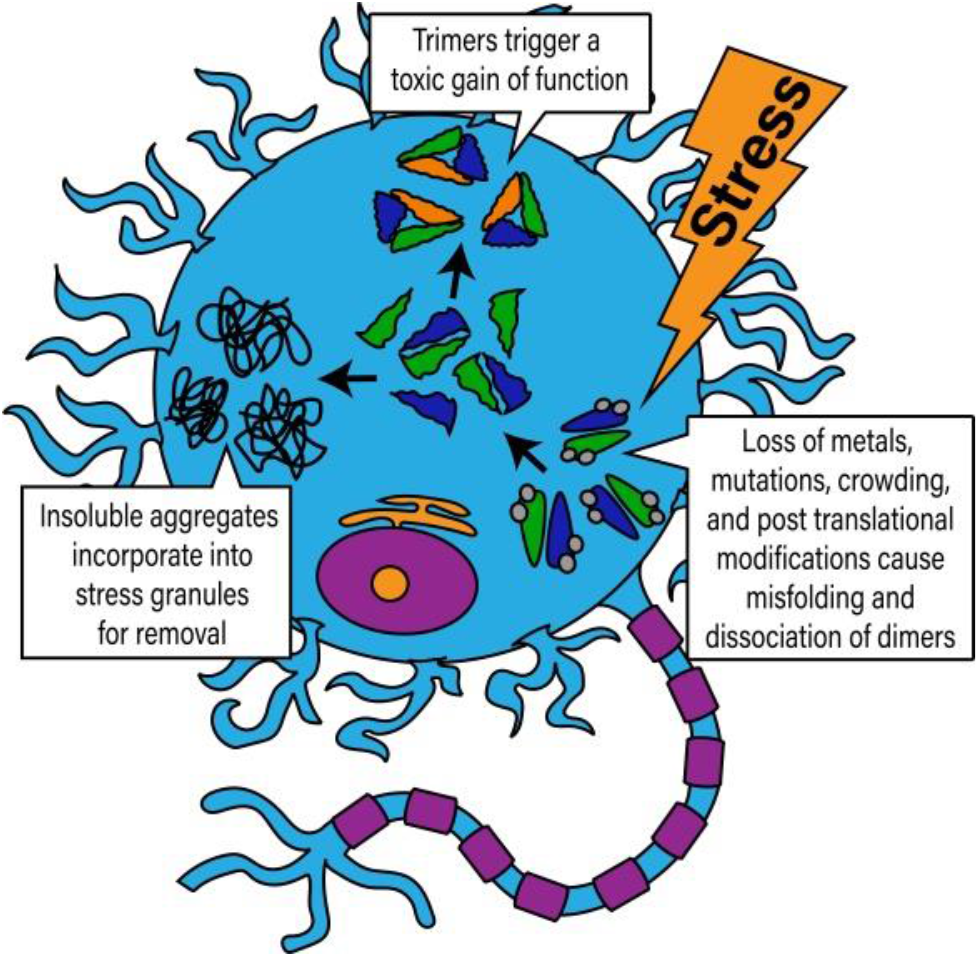
The formation of fibrils is a natural protective mechanism in cells during stress. Formation of off-pathway oligomers is a toxic by product contributing to cell death in multiple diseases.

We corroborate Proctor and Zhu’s findings that formation of fibrils rescues cells while stabilizing the trimeric form of SOD1 increases cytotoxicity (31) (32). Our super stable trimer mutants of SOD1 may be used in further studies to determine the trimer’s exact structure and protein-protein interactions. Our concentration-based correlation method for determining if an intermediate conformation is on- or off-pathway has the potential to be applied to other aggregating proteins with toxic intermediates where the intermediates cannot be characterized by ThT, such as amyloid beta. Finally, the initial structural information we achieved through CD and limited proteolysis shows how structurally different the trimer is from other conformations of SOD1. These drastic differences in structure explain why most misfolded SOD1 antibodies do not recognize trimers, while those that do bind to trimer (C4F6) do not bind to amyloid fibrils or native dimers (8) (31) (51). This challenge has led to the split opinion on the extent of SOD1’s role in sporadic ALS. Studies that use antibodies that identify trimeric SOD1 find misfolded SOD1 in both sALS and fALS, while studies that utilize antibodies that do not identify SOD1 trimers do not observe the same misfolded accumulation in sALS (52) (53) (54). The evidence for SOD1 trimers in sALS patients suggests there is an overarching effect of trimeric SOD1 throughout ALS opposed to solely in SOD1 fALS cases.

Different therapeutics targeting SOD1 in ALS have recently focused on stabilizing the native dimers (55) (56) (57) (58) (59). While this strategy is promising it is not likely to work on later stage patients who already have accumulated misfolded SOD1, and currently there is no universal biomarker to screen for ALS before symptoms begin. Another strategy is reducing the overall amount of SOD1, Tofersen is an antisense oligonucleotide for SOD1 which aims to reduce SOD1 transcription and it has already shown promise in clinical trials (60). Our findings also open the door to new therapeutics for ALS focusing on promoting fibril formation (which has recently been investigated in Alzheimer’s Disease (61)), destabilizing the trimer directly, or determining and preventing the downstream toxic interactions with the trimer. Studying the toxic role of smaller soluble oligomers in neurodegenerative diseases is gaining momentum, showing that the toxic trimeric form of SOD1 is off-pathway leads the way for new therapeutic and biomarker development.

## Methods

### Purification of SOD1

SOD1 WT and mutants are first overexpressed in the SOD1 knockout yeast line EG118 then purified using ammonium sulfate precipitation followed by hydrophobic interaction chromatography and size exclusion chromatography as previously described (62) (5). Freshly purified samples are stored at -20°C until use.

### Thioflavin T

We dilute SOD1 protein samples to 300 μM in 20 mM Tris 150 mM NaCl pH 7.4, the concentration is determined using UV-vis spectrophotometry absorbance at 280 nm with an extinction coefficient of 10800 M^-1^cm^-1^. All samples are diluted to 300 μM and tris(2-carboxyethyl)phosphine is added to a final concentration of 30 mM and the samples are incubated for 1 hour at 37°C with 300 rpm shaking to reduce any aggregated protein to apo-monomers. After incubating, we dilute each sample 1:10 to achieve a final protein concentration of 30 μM. We then immediately add ThT to 40 μM and pipette 200 μL per well in a 96 well black wall clear bottom plate (three wells per construct). We add a 3 mm glass bead to each well (63), then seal the plate with an adhesive lid to prevent evaporation. We use a SpectraMax i3 microplate detection system (Molecular Devices) to measure the kinetics of aggregation. The machine is pre-warmed to 37°C for 30 minutes, excitation and emission filters are set to 440 nm and 480 nm respectively, fluorescence measurements are taken from the bottom of the plate at 2.5 minute intervals for 144 hours, and the plate is orbitally shaken between readings.

### NSC-34 Cell Growth and Lysis

We grow neuroblastoma spinal cord hybrid cells (NSC-34) in 50:50 DMEM: F12 containing 10% (vol/vol) FBS and 1% penicillin/ streptomycin. We plate ∼7,000 NSC-34 cells per well in a 12 well tissue culture plate or ∼20,000 cells per well in a 6 well tissue culture plate. The cells are differentiated for at least two days by adding retinoic acid to a concentration of 10 μM, we transfect the cells with either WT or mutant SOD1 pCI-Neo plasmid using JetOptimus transfection reagent (VWR) according to the manufacturer’s instructions (31). Transfection efficiency is confirmed using a co-expressed mCherry protein. Any time the media is removed post-transfection, any floating cells are saved in cold 1X PBS. Three days after transfection the cells are harvested using 0.25% Trypsin (Gibco) to remove them from the wells, these cells are combined with the floating cells and resuspended in either cold Lactate Dehydrogenase Activity (LDH) reaction buffer or 1X PBS with EDTA free protease inhibitor (Protease Inhibitor for Mammalian Cells, Dot Scientific). Five 3 mm glass beads are added to each vial before mixing ten times on a vortex for ten seconds each time to lyse the cells; cells are placed on ice in between mixing.

### SOD1 Inclusion formation in NSC-34 Cells

NSC-34 lysate from two wells of a 6 well plate is used for each sample. All samples are centrifuged for 15 minutes at 13,000 x g to pellet the insoluble fraction (containing any insoluble aggregates). The supernatant is removed and saved for immunoblots while the pellet is resuspended in 200 μL of 1X PBS and then centrifuged again. This process is repeated two more times to wash the pellets of any soluble protein. The remaining insoluble pellet is resuspended in PBS with reducing solution (0.4 M Tris, 60% glycerol, 120 mg/mL SDS, and 93 mg/mL DTT) then heated to 95°C for 15 minutes to reduce the protein to monomers. A bicinchoninic acid (BCA) assay kit (Thermo Fisher) is used to standardize the samples to the same protein level, then the samples are diluted 1:10, 1:50, and 1:100 and 2 μL of each sample are dotted on activated 0.2 μm PVDF membrane in duplicate. The blots are allowed to air dry for 1.5 hours before they are blocked with 5% BSA solution in 1X TBST for one hour, we then incubate the blots with primary antibody (anti-SOD1, Sigma SAB5200083, 1:1000) overnight at 4°C. The following day the blots are incubated with secondary antibody (anti-Rabbit HRP, Life Tech A16096, 1:1000) for one hour at room temperature. Pierce ECL substrate is used to visualize the HRP; the blots are imaged using a Chemidoc Imager then quantified using ImageJ.

### Lactate Dehydrogenase Assay

Lysed cells from three wells of a 12 well plate are used for each sample. The assay is performed following the manufacturer’s instructions (BioVision Lactate Dehydrogenase Activity Colorimetric Assay Kit).

### Immunoblots

NSC-34 lysate from two wells of a 6 well plate are used for each sample. All samples are centrifuged for 15 minutes at 13,000 x g to pellet the lysed cells. We denature the lysates and resolve using 10% (vol/vol) SDS/PAGE before blotting on PVDF membrane using a TransBlot Turbo (BioRad). Blots are incubated with antibodies against SOD1 (Sigma SAB5200083) or Actin (MP Biomedicals clone C4) followed by secondary antibody incubation. We visualize the blots using a ChemiDoc MP system (BioRad).

### SOD1 Aggregation Time Course Experiments

We dilute purified de-metallated SOD1 to 30 μM in 20 mM Tris 150 mM NaCl pH 7.4. The dilute sample is divided in half, half the sample is frozen while the other half is rotated at 37°C. Initially, and at 30 minute increments, we remove 100 μL from the sample and run SEC on a Superdex S200 column (the column was run for at least 30 minutes prior to the first time point). After 12 hours we run a zero time point on the frozen half and begin incubating it. Time points are run every 30 minutes, alternating samples, for at least 60 hours. We calculate the area under the curve of the chromatograph for each time point using the Unicorn 7 software. The total area at each time point is subtracted from the highest total area to account for any protein lost to insoluble fibrils (insoluble protein cannot be measured by the column). The correlation coefficient, coefficient of determination, and P value are calculated using Python.

### Circular Dichroism

We de-metallate samples of SOD1 mutants by dialyzing each against a 50 mM Sodium Acetate and 10 mM EDTA solution at pH 3.8 for 1.5 hours to obtain apo-monomer, trimer, and larger aggregates. The samples are then dialyzed against a 20 mM Tris pH 7.4 solution twice overnight to return to a neutral pH before isolating the different size aggregates by SEC using a 25 mL Superdex 200 column. The fractions for each size aggregate are pooled and concentrated to at least 0.1 mg/mL using an Amicon Ultra filter (Millipore) with a 10 kDa filter cutoff. We determine the concentrations using a BCA assay kit. We then determine the secondary structure characteristics of each sample using a Jasco J-1500 circular dichroism spectrophotometer. We place each sample in a 0.1 mm path-length (200 uL) cuvette, and record the spectra range at 185-240 nm with 0.5 nm data pitch at a scanning speed of 50 nm/min for three times and then take the average. The fractional content of the helix, sheet, turn, and other structural elements is determined using “BESTSEL” to analyze the experimental data.

### Limited Proteolysis

We isolate insoluble fibrils for limited proteolysis by incubating SOD1 samples in 20 mM Tris 150 mM NaCl pH 7.4 buffer for one week at 37°C, then we centrifuge the samples at 15,000 g for 15 minutes. We wash the pelleted insoluble fibrils twice with the aggregation buffer then resuspend the pellet and determine the concentration using BCA. We isolate SOD1 trimers by SEC using a Superdex 200 column. We digest the samples using 82 μM pepsin with 10 μM protein in 100 mM sodium acetate pH 3.5 for multiple time points from 1 minute to 30 minutes. The digestions are quenched by adjusting the pH back to neutral using 1M Tris pH 8 and adding PMSF to 1 mM before flash freezing. We analyze the digested samples with Mass Spectrometry using the Sciex 5600 triple tof (AB SCIEX) and the software Byonic (Penn State College of Medicine Mass Spectrometry and Proteomics Core Facility, RRID:SCR_017831).

## Acknowledgments

We thank Esther Choi, Madison Kuhn, Jiaxing Chen, Joshua Reynolds, and Dr. Deepkamal Karelia for valuable discussions. We thank the Mass Spectrometry & Proteomics College of Medicine Core as well as the X-Ray Crystallography Core for their assistance with experiments. We acknowledge support from the National Institutes for Health (R35 GM134864 to N.V.D.) and the Passan Foundation.

