## Supplementary Information for "Toxic SOD1 trimers are off-pathway in the formation of amyloid-like fibrils in ALS"

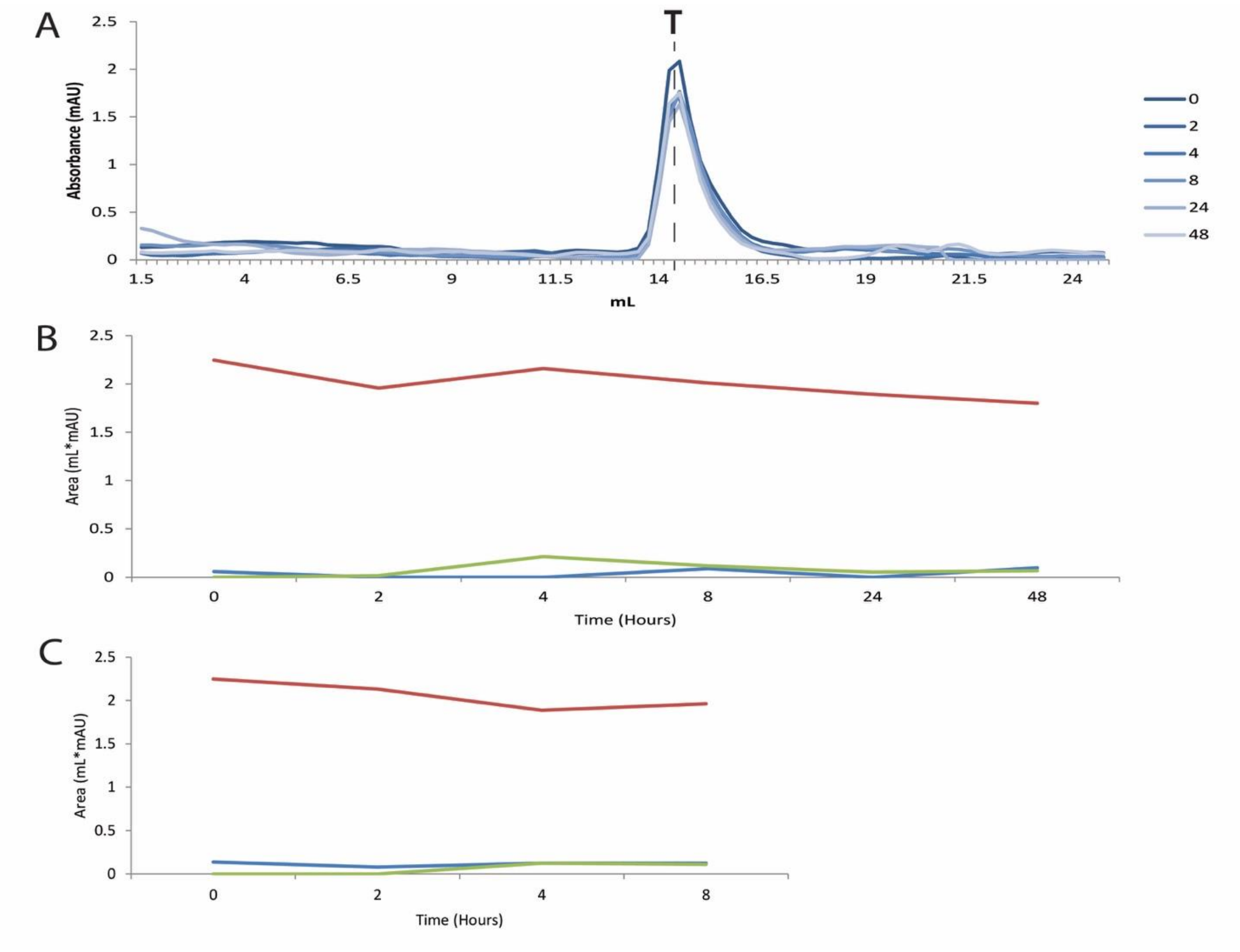


**Fig. S1.** Using size exclusion chromatography, we showed that the stabilized trimers remain in the trimeric form for at least 8 hours when incubated at 37°C. (A) Double mutant H46Q-G108H, beginning from a population of all trimers, remained as trimers across 48 hours (darkest= 0 hours, lightest =48 hours). Less trimer stable constructs (such as F20L) had increased populations of larger oligomers and monomers. (B) We quantify the peak areas during size exclusion chromatography and further show that the SOD1 trimers (red) stay stable and there are very few fluctuations in larger oligomers/ insoluble aggregates (green) and monomers (blue). (C) Similarly F20L-H46Q also stays stable as trimers for 8 hours.

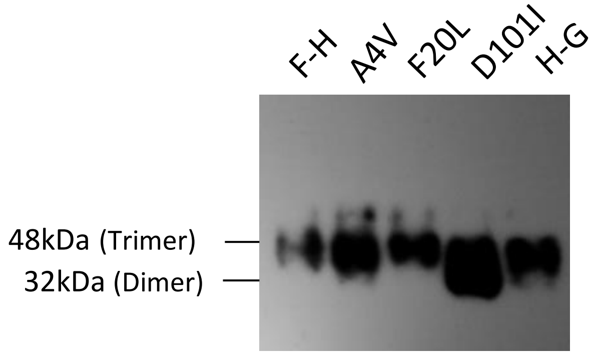


**Fig. S2.** We performed Western blotting using an anti-misfolded SOD1 antibody (anti-C4F6, Medimabs) and show that the trimer stabilizing mutants are forming trimers within NSC-34 cells. Anti-C4F6 is one of the few misfolded SOD1 antibodies which has been previously shown to recognize trimers. All of the trimer stabilizing mutants (F20L-H46Q [F-H], F20L, and H46Q-G108H [H-G]) as well as A4V have a distinct trimer band at 48kDa. D101I, which is designed to destabilize trimers, has a trimer band along with a larger lower molecular weight band which is the correct size for dimeric SOD1 (32kDa) this dimer band might be misfolded which is why it is appearing with the misfolded SOD1 antibody.

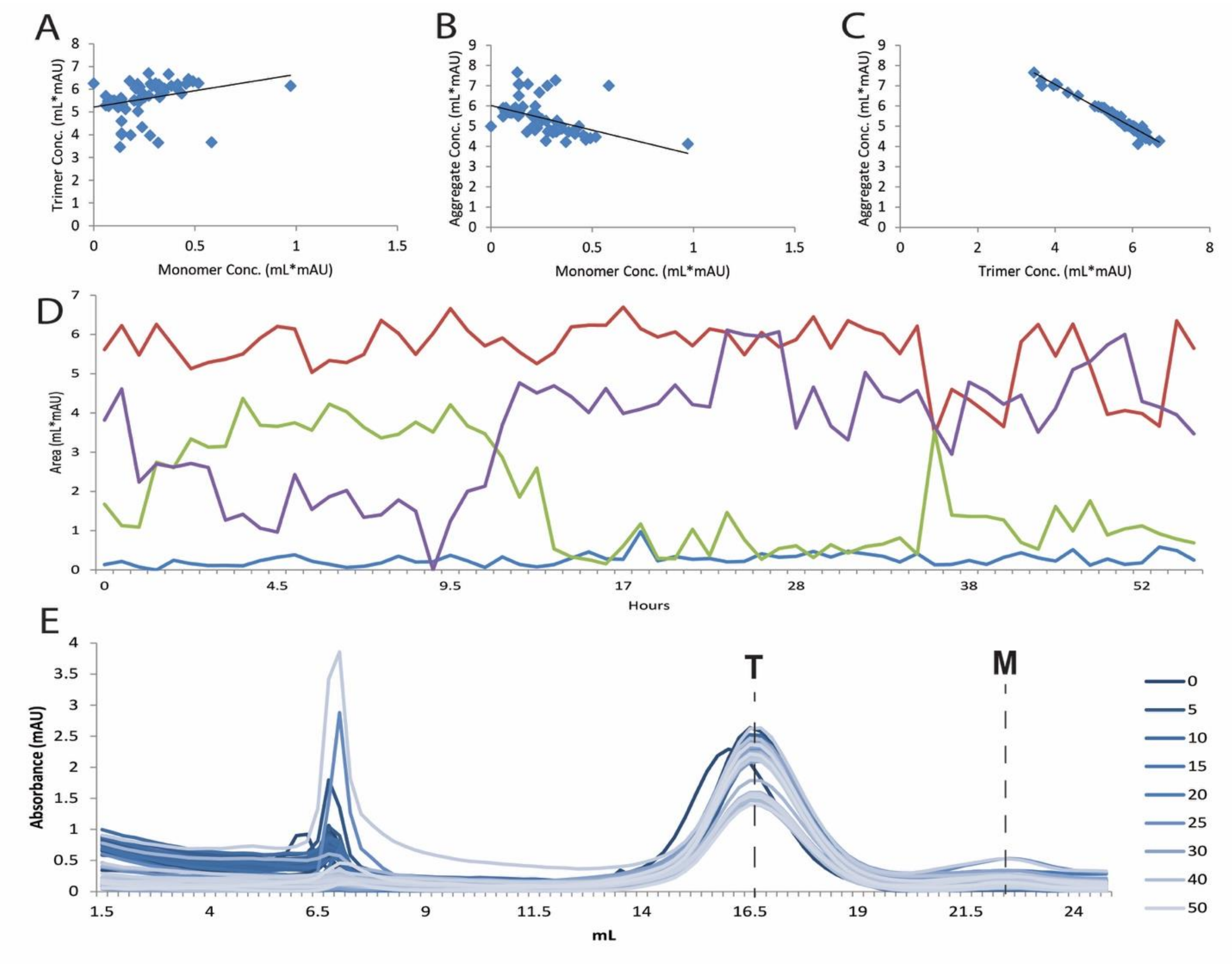


**Fig. S3.** Size exclusion chromatography time course for A4V 1 (n=64).
(A) The monomer and trimer concentrations have a correlation coefficient of 0.29 [$p=1.97x10^{-2}$]. (B) The monomer and aggregate (insoluble and larger soluble) concentrations have a correlation coefficient of -0.46 [$p=10^{-5}$]. (C) The trimer and aggregate concentrations have a strong negative correlation coefficient of -0.98 [$p=6.93x10^{-48}$]. (D) Quantification of the peak areas throughout the time course shows fluctuations between the different species (trimer-red, monomer- blue, soluble larger oligomer- green, insoluble aggregates- purple) that are not seen when further stabilizing the trimer. (E) The size exclusion chromatograph shows the change in different size aggregates across 50 hours (darkest= 0 hours, lightest =50 hours), the area under these curves was quantified to determine the correlation coefficients.

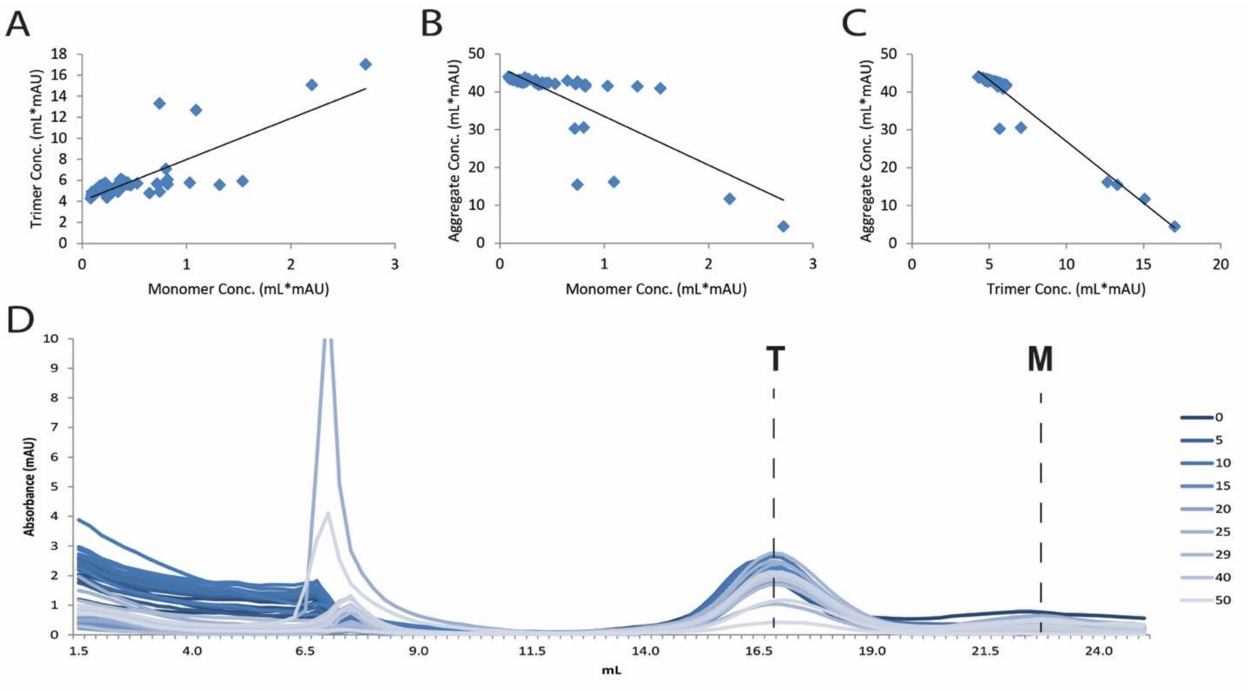


**Fig. S4.** Size exclusion chromatography time course for A4V 2 (n=51).
(A) The monomer and trimer concentrations have a correlation coefficient of 0.78 [$p=1.9x10^{-11}$]. (B) The monomer and aggregate (insoluble and larger soluble) concentrations have a correlation coefficient of -0.75 [$p=2.17x10^{-10}$]. (C) The trimer and aggregate concentrations have a strong negative correlation coefficient of -0.92 [$p=7.34x10^{-22}$]. (D) The size exclusion chromatograph shows the change in different size aggregates across 50 hours (darkest= 0 hours, lightest =50 hours), the area under these curves was quantified to determine the correlation coefficients.

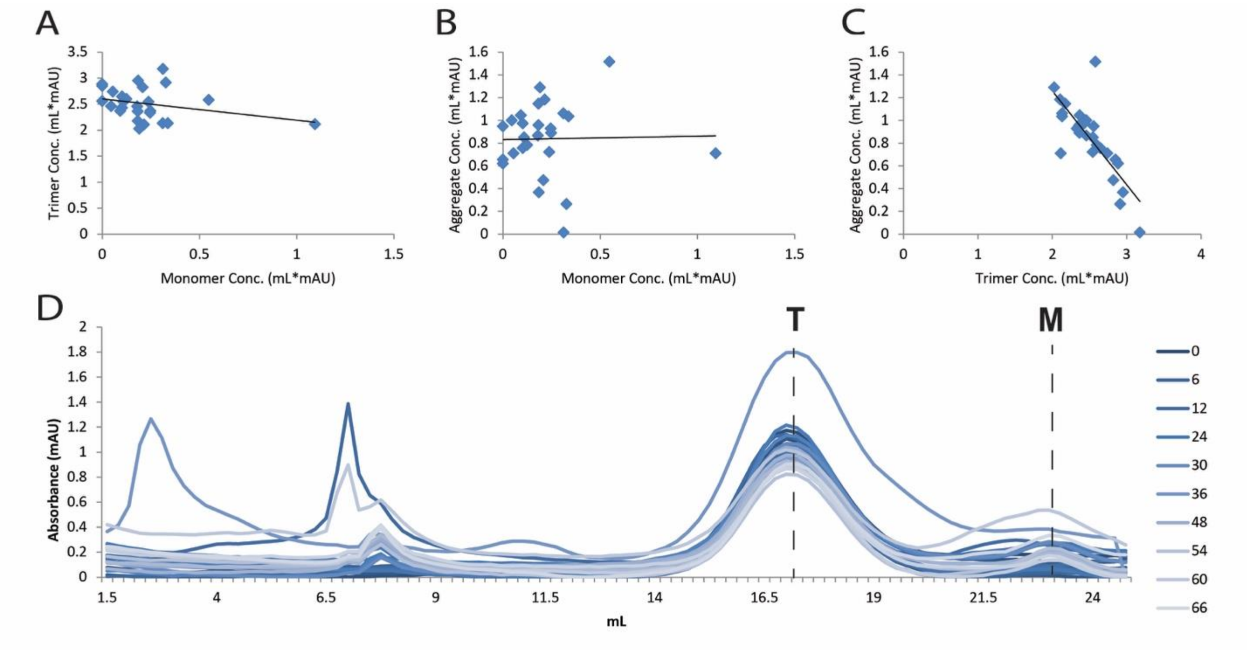


**Fig. S5.** Size exclusion chromatography time course for A4V 3 (n=26).
(A) The monomer and trimer concentrations have a correlation coefficient of -0.29 [p=0.144]. (B) The monomer and aggregate (insoluble and larger soluble) concentrations have a correlation coefficient of 0.02 [p=0.811]. (C) The trimer and aggregate concentrations have a strong negative correlation coefficient of -0.78 [$p=2.58\times10^{-5}$]. (D) The size exclusion chromatograph shows the change in different size aggregates across 50 hours (darkest= 0 hours, lightest =50 hours), the area under these curves was quantified to determine the correlation coefficients.

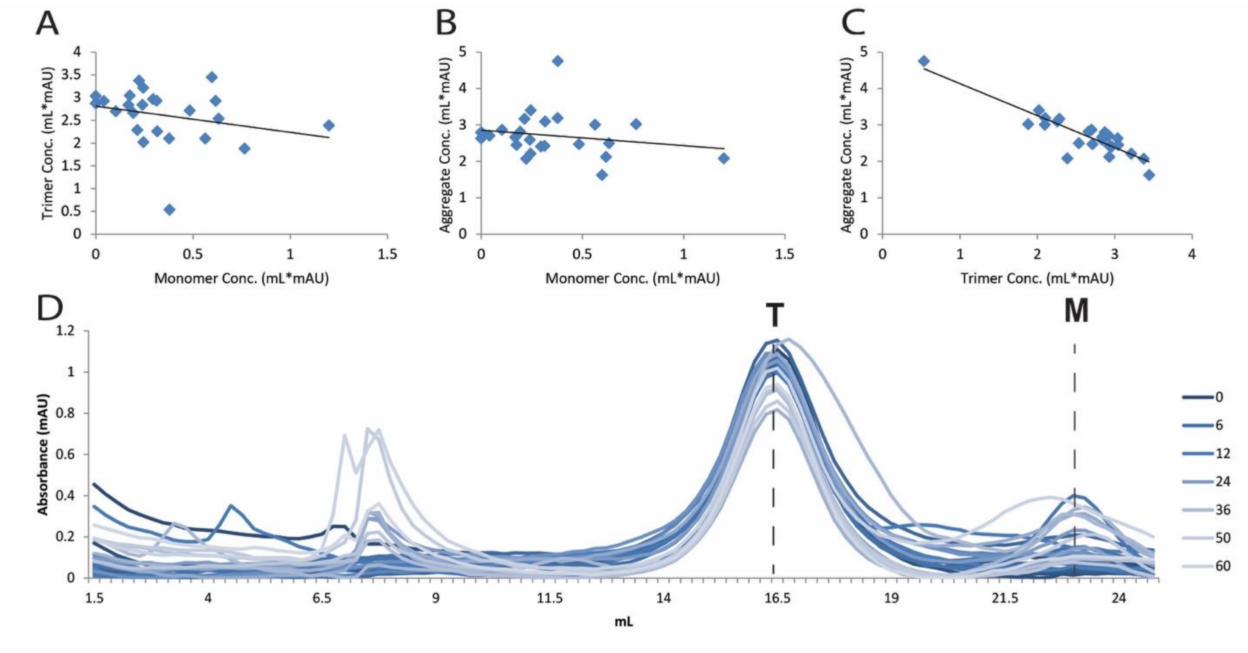


**Fig. S6.** Size exclusion chromatography time course for F20L 1 (n=25).
(A) The monomer and trimer concentrations have a correlation coefficient of -0.27 [p=0.20]. (B) The monomer and aggregate (insoluble and larger soluble) concentrations have a correlation coefficient of -0.20 [p=0.339]. (C) The trimer and aggregate concentrations have a strong negative correlation coefficient of -0.89 [$p=2.16\times10^{-9}$]. (D) The size exclusion chromatograph shows the change in different size aggregates across 50 hours (darkest= 0 hours, lightest =50 hours), the area under these curves was quantified to determine the correlation coefficients.

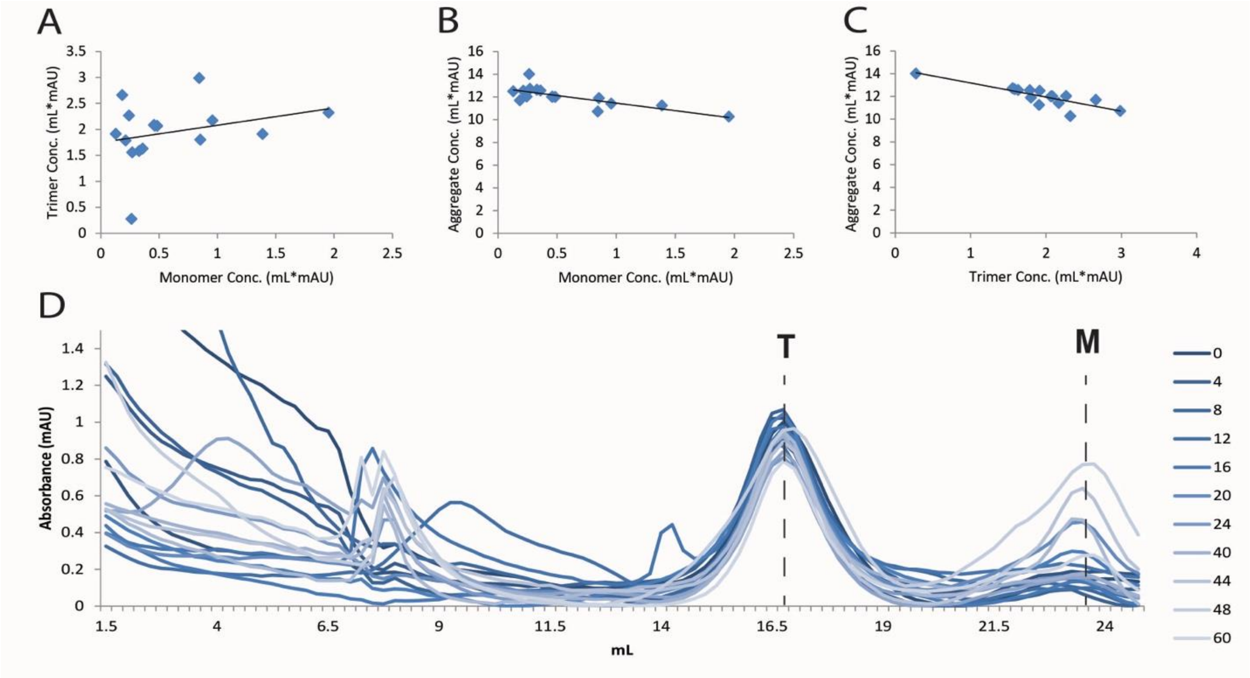


**Fig. S7.** Size exclusion chromatography time course for F20L 2 (n=16).
(A) The monomer and trimer concentrations have a correlation coefficient of 0.285 [p=0.29]. (B) The monomer and aggregate (insoluble and larger soluble) concentrations have a correlation coefficient of -0.77 [$p=5.5x10^{-4}$]. (C) The trimer and aggregate concentrations have a strong negative correlation coefficient of -0.83 [$p=5.77x10^{-5}$]. (D) The size exclusion chromatograph shows the change in different size aggregates across 50 hours (darkest= 0 hours, lightest =50 hours), the area under these curves was quantified to determine the correlation coefficients.
